# Restoring neurovascular coupling in Alzheimer’s disease tauopathy through M1 mAChR modulation

**DOI:** 10.64898/2026.08.18.745579

**Authors:** Wesam Bassiouni, Mohamed Abdelnaby, En-Hung Ai, Khaled S. Abd-Elrahman

## Abstract

Alzheimer’s disease is characterized by progressive cognitive decline and early cerebrovascular dysfunction, including impaired neurovascular coupling (NVC) and reduced cerebral blood flow (CBF). Tau pathology is a major driver of these deficits, yet therapeutic strategies targeting tau-induced neurovascular dysfunction remain limited. The M1 muscarinic acetylcholine receptor (M1 mAChR) is a promising therapeutic target because of its critical role in cognition. We previously demonstrated that pharmacological activation of M1 mAChR improves cognitive function and neuronal survival in amyloid-based Alzheimer’s disease mouse models through sex-specific mechanisms. However, whether M1 mAChR activation restores tau-mediated NVC deficits remains unknown. P301S mice were used as a model of tauopathy. Cognitive function was evaluated using the novel object recognition and Morris water maze tests, and NVC was assessed by measuring whisker stimulation-induced changes in CBF using laser speckle contrast imaging. Following baseline measurements, mice received an acute intraperitoneal injection of VU0486846, a selective M1 mAChR positive allosteric modulator (3 mg/kg), and CBF responses were reassessed over time. P301S tau mice exhibited impaired recognition and spatial memory functions, associated with reduced whisker stimulation-induced increase in CBF, indicative of impaired NVC response, while acute treatment with VU0486846 reversed these changes in NVC. This rescuing effect of VU0486846 was observed earlier in female tau mice compared to males, suggesting a sex-biased effect of M1 mAChR modulation. These findings demonstrate that M1 mAChR positive allosteric modulation reverses tau-induced neurovascular dysfunction, supporting M1 mAChR activation as a promising disease-modifying approach for Alzheimer’s disease. The earlier improvement observed in females further suggests that therapeutic efficacy is influenced by biological sex.

**Abbreviated Summary:** Bassiouni et al. report that tauopathy impairs neurovascular coupling in mice. Acute positive allosteric modulation of the M1 muscarinic acetylcholine receptor selectively restored neurovascular coupling in tau mice. These findings identify M1 receptor modulation as a promising therapeutic strategy to restore neurovascular function in Alzheimer’s disease.

## Introduction

Alzheimer’s disease is a progressive neurodegenerative disorder marked by cognitive decline and neuronal loss.^1^ Key pathological hallmarks of Alzheimer’s disease include the accumulation of β-amyloid plaques and hyperphosphorylation of the microtubule-associated protein tau, both of which contribute to neurotoxicity and disease progression.^2^ Alzheimer’s disease is also characterized by early cerebrovascular dysfunction including impaired neurovascular coupling (NVC), a tightly regulated process by which increases in neuronal activity trigger corresponding increases in local cerebral blood flow (CBF) to meet metabolic demands, a mechanism mediated by coordinated interactions among neurons, astrocytes and vascular cells.^3^ In Alzheimer’s disease, this process becomes disrupted, leading to impaired delivery of oxygen and glucose to active brain regions and contributing to synaptic dysfunction and cognitive decline. Interestingly, this reduction in CBF and impaired NVC occur during the early stages of the disease and is highly correlated with cognitive decline.^4^ Previous reports show that tau pathology plays a direct role in this dysfunction, as hyperphosphorylated tau can impair astroglial signaling, promote neuroinflammation, and induce endothelial dysfunction, all of which compromise vascular responsiveness to changes in metabolic demand.^5, 6^

The cholinergic system is one of the earliest neurotransmitter systems affected during Alzheimer’s disease progression. Cholinergic neurons in the basal forebrain are susceptible to neurodegenerative changes, leading to impaired neurotransmission.^7, 8^ Among cholinergic receptors, the M1 muscarinic acetylcholine receptor (M1 mAChR) is the most abundant G protein–coupled receptor subtype in the hippocampus and cortex, and plays a crucial role in regulating cognitive processes, neuronal survival, and synaptic plasticity.^9, 10^ Our previous work identified the M1 mAChR as a promising therapeutic target for Alzheimer’s disease. Indeed, we demonstrated that VU0486846, a novel orally bioavailable M1 mAChR positive allosteric modulator (PAM), improves cognitive function in Alzheimer’s disease mouse models while reducing β-amyloid pathology, preventing neuronal loss, and attenuating reactive astrogliosis.^8, 11, 12^ Given the central role of astrocytes in regulating NVC ^13^ and the mounting evidence that astrocytes express M1 mAChRs,^14^ these findings suggest that M1 mAChR activation may regulate NVC in addition to its well-established effects on neuronal function. This hypothesis is further supported by clinical evidence demonstrating that enhancing cholinergic signaling with acetylcholinesterase inhibitors improves CBF in patients with Alzheimer’s disease, although the specific muscarinic receptor subtype mediating these effects remains unknown.^15, 16^ Together, these findings provide a strong rationale for investigating whether M1 mAChR activation can preserve neurovascular function and reverse tau-induced neurovascular dysfunction in Alzheimer’s disease.

Here we report that male and female P301S tau mice exhibit comparable cognitive impairment and deficits in NVC compared with wild-type controls. Interestingly, VU0486846 significantly restored NVC in P301S tau mice of both sexes with no noticeable difference in wild-type controls, indicating a disease-specific effect. Our data also demonstrate that female P301S tau mice exhibited a faster NVC response following VU0486846 treatment than male mice, indicating more rapid restoration of neurovascular dynamics in females. These findings identify M1 mAChR positive allosteric modulation as a promising strategy to restore neurovascular function in Alzheimer’s disease and support further investigation of M1 mAChR-targeted therapies for Alzheimer’s disease and related tauopathies.

## Materials and Methods

### Reagents

VU0486846 ((R)-4-(4-(1H-Pyrazol-1-yl) benzyl)-N-((1S,2S)-2-hydroxycyclohexyl)-3,4 dihydro-2H-benzo [b] [1,4] oxazine-2-carboxamide) (Cat# 3271) was obtained from Axon Medchem (Virginia, USA).

### Animals

All animal experimental protocols were approved by the University of British Columbia Animal Care Committee (Protocol ID: A23-0045) and in accordance with the Canadian Council of Animal Care (CCAC) guidelines. Mice were obtained from The Jackson Laboratories and bred to establish littermate-controlled male and female wild-type and P301S tau mice (B6; C3-Tg (Prnp-MAPT*P301S) PS19Vle/J, Strain#:008169, RRID: IMSR_JAX:008169). Mice were group-housed in cages of 2 or more animals, received food and water ad libitum and maintained on a 12-h light/12-h dark cycle at 24°C. Wild-type and P301S tau mice were aged to 10 months before starting the behavioral experiments and measurement of CBF.

### Morris water maze

To test the mice spatial memory, the Morris water maze test was performed using a plastic pool (diameter: 120 cm and depth: 50 cm), filled with water that was made opaque by addition of white nontoxic water-soluble paint. Temperature was maintained at 25°C throughout the whole experiment to prevent hypothermia. An escape platform (10 cm in diameter) was placed 25 cm from the perimeter and submerged 1 cm under the water surface. Visual cues included a ‘X’ and square were placed on the front and right walls of the maze room opposite to the target quadrant as spatial references. Mice were trained for 4 days (four trials per day, 60 s each and 20 min between trials) to find the submerged platform at a fixed position from a random starting point of one of the four quadrants and the starting quadrant was changed on each trial day. The trial ends either when it reaches 60 s or when the mice find the platform and stay on it for 5 s whatever comes first. If the mice failed to find the platform within 60 s, they were guided to the platform by the experimenter. The escape latency, which is calculated as the time needed to reach the platform in (s), was measured using ANY-Maze tracking software connected to an automated video tracking camera, and the average of the four daily trials was plotted over the testing days. On day 5, known as the probe trial which is a single trial of 60 s, the platform was removed and mice were allowed to swim freely in the pool and the time in (s) spent in the target quadrant was measured.^17, 18^

### Novel object recognition

To assess the mice non-spatial recognition memory, the novel object recognition test was performed on a two-day experiment. In the first day, mice were placed in an empty box (45 × 45 × 45 cm), habituated for 10 min and were then returned to their home cage. One hour later, two identical objects were placed in the box 5 cm from the edge and 5 cm apart and mice were returned to the box and allowed to explore the two objects for 5 min. The time spent exploring each object was recorded using ANY-Maze tracking software connected to an automated video tracking camera. Mice were considered to be exploring the objects if their snouts were within 1 cm of the object. 24 hours later, one of the objects was replaced with a novel object and the experiment was repeated for each animal. Data were interpreted using a discrimination index as the time spent exploring the familiar object subtracted from the time spent exploring the novel object divided by the total time spent exploring both objects. The discrimination index is calculated as a fraction from -1 to 1, where a positive value indicates that mice are spending more time exploring the novel object, indicating preserved memory function, while a negative value means that they are spending more time exploring the familiar one, indicating impaired memory.^17, 19^

### Laser speckle contrast imaging and VU0486846 treatment

To measure the changes in NVC, mice were anesthetized using isoflurane and fixed in a stereotaxic apparatus, and anesthesia was maintained under 2% isoflurane for the whole experiment. Following complete non-responsiveness to external stimuli, the scalp was surgically opened and the skull exposed. Changes in baseline CBF were recorded using laser speckle contrast imaging LSCI (RFLSI III, RWD Life Sciences). Whisker stimulation was performed on the right side of anesthetized mouse using a manually controlled electric brush for 20 s to stimulate the brain activity and cerebral perfusion as a measurement of NVC response. For each mouse, three technical trials were acquired with one min gap in between, from three different regions of interest (ROIs). The change in CBF was recorded for each trial over a duration of one min (20 s before stimulation, 20 s stimulation, 20 s after stimulation), calculated as a percentage change in CBF (%Δ in CBF) compared to the baseline before stimulation, and averaged among the different trials and different ROIs.^20^

Following baseline CBF recording, mice received an an acute intraperitoneal injection of VU0486846 (3 mg/kg) followed by reassessment of changes in CBF every 15 min for 60 min and %Δ in CBF was calculated and compared among the different groups. The slope of whisker stimulation–induced increases and stimulation-offset–induced decreases in CBF was quantified using linear regression analysis applied to the time interval beginning at the onset of the response and ending upon reaching a plateau to obtain the slope to plateau, and from the onset of decline till reaching the baseline to obtain the slope to baseline.

### Data analysis

Data are expressed as the mean ± standard error of the mean (SEM) of n independent experiments (where n represents an individual mouse). For two groups comparison, unpaired student t-test was used. For multiple comparisons, two-way analysis of variance (ANOVA) followed by Fisher’s LSD test was used (GraphPad Prism version 10, California, USA). Statistical significance was considered at p < 0.05.

## Results

### P301S tau mice exhibit impaired cognitive function

To first confirm that P301S tau mice exhibited cognitive deficits at this age, we assessed both non-spatial recognition memory and spatial learning and memory using the novel object recognition and Morris water maze tests, respectively. Novel object recognition test showed that both female and male P301S tau mice exhibited impaired recognition memory as evidenced by their lower discrimination indices compared to their sex-matched wild-type controls, reflecting their declined ability to discriminate between novel and familiar objects, while wild-type controls were functionally able to distinguish between the two objects as evidenced by their higher discrimination indices (**Fig. 1A**). Using Morris water maze, both female and male P301S tau mice showed impaired spatial memory as evidenced by having longer escape latency and shorter time spent in target quadrant compared to their sex-matched wild-type controls which showed improved escape latency across the training days of the trails and longer time spent in target quadrant in the probe day (**Fig. 1B &C**), reflecting their preserved learning and memory functions. These findings are consistent with previous reports of cognitive impairment in P301S mice at earlier disease stages.^21–23^

**Figure 1.**
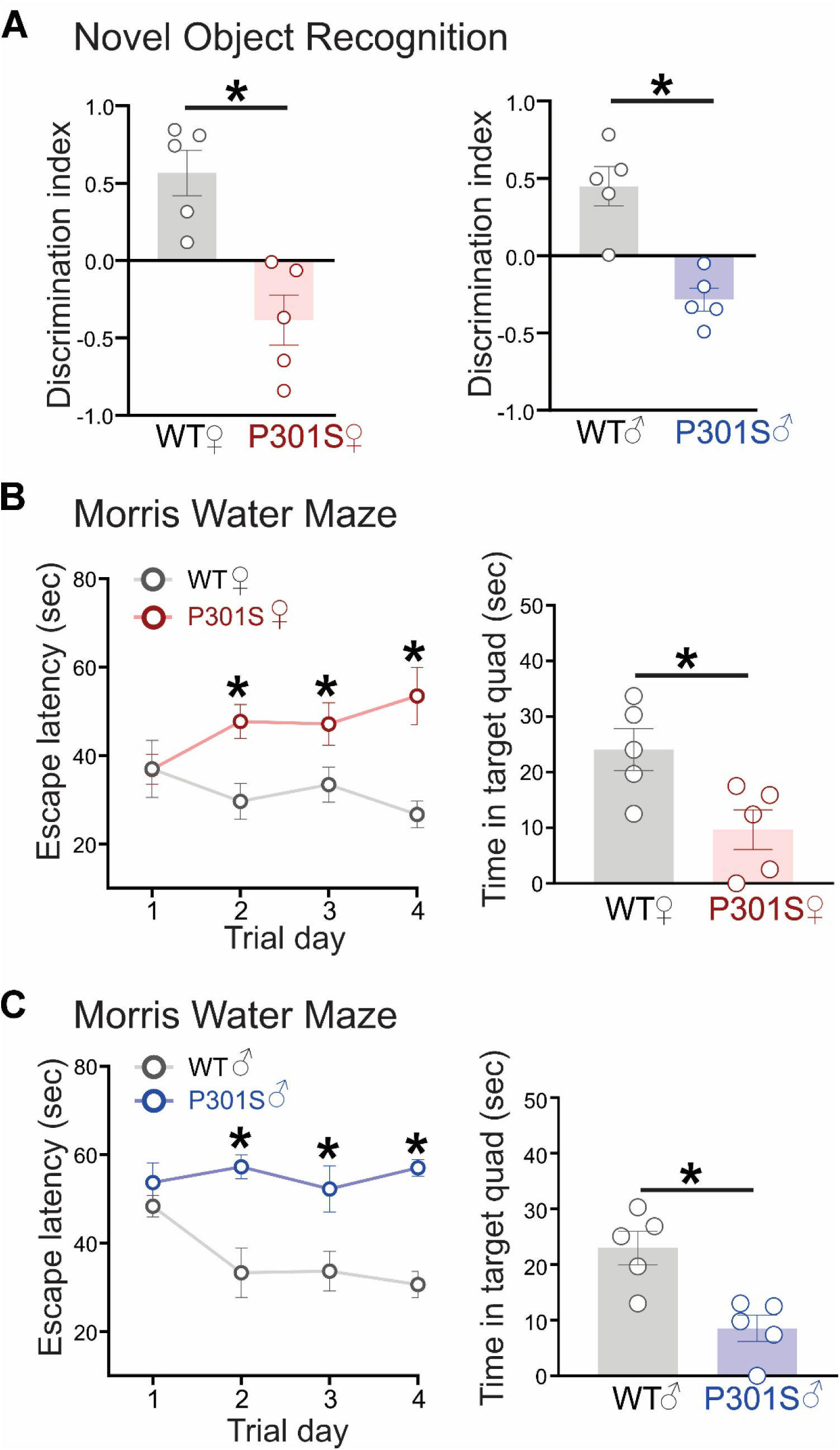
Impaired cognitive function in male and female P301S tau mice. (**A**) **Novel object recognition:** Discrimination index between novel and familiar objects in 10-month-old female (left panel) and male (right panel) wild-type (WT) and P301S tau mice. (**B&C**) **Morris water maze:** Escape latency (left panel) and time spent in target quadrant (right panel) in (**B**) female and (**C**) male WT and P301S tau mice. *p<0.05 by unpaired student t-test or two-way ANOVA (n=5 per group). Error bars denote SEM.

### P301S tau mice exhibit deficient neurovascular coupling responses

We next examined whether the cognitive deficits observed in P301S tau mice at this age are accompanied by impaired NVC. To assess neurovascular responses, changes in CBF were measured using laser speckle contrast imaging during whisker stimulation, which evokes neuronal activity and the corresponding increase in local cerebral perfusion.^20, 24^ NVC responses were then compared between male and female P301S tau mice and their age-matched wild-type controls. Impaired NVC was observed in both female and male P301S tau mice compared to their sex-matched wild-type controls as evidenced by the time course reduction in %Δ in CBF upon whisker stimulation (**Fig. 2A &D**) as well as the reduced peak %Δ in CBF (**Fig. 2C &F**) observed in P301S tau mice compared to controls, confirming that the cognitive decline observed in P301S tau mice is associated with reduced NVC. No significant differences in basal CBF acquired before whisker stimulation were observed between wild-type and P301S tau mice of both sexes (**Fig. 2B &E**). These findings suggest that impaired NVC, rather than altered basal cerebral perfusion, is closely associated with cognitive deficits in P301S tau mice, underscoring the importance of targeting neurovascular dysfunction as a therapeutic strategy for tauopathies and establishing the P301S model as a valuable platform for evaluating such interventions.

**Figure 2.**
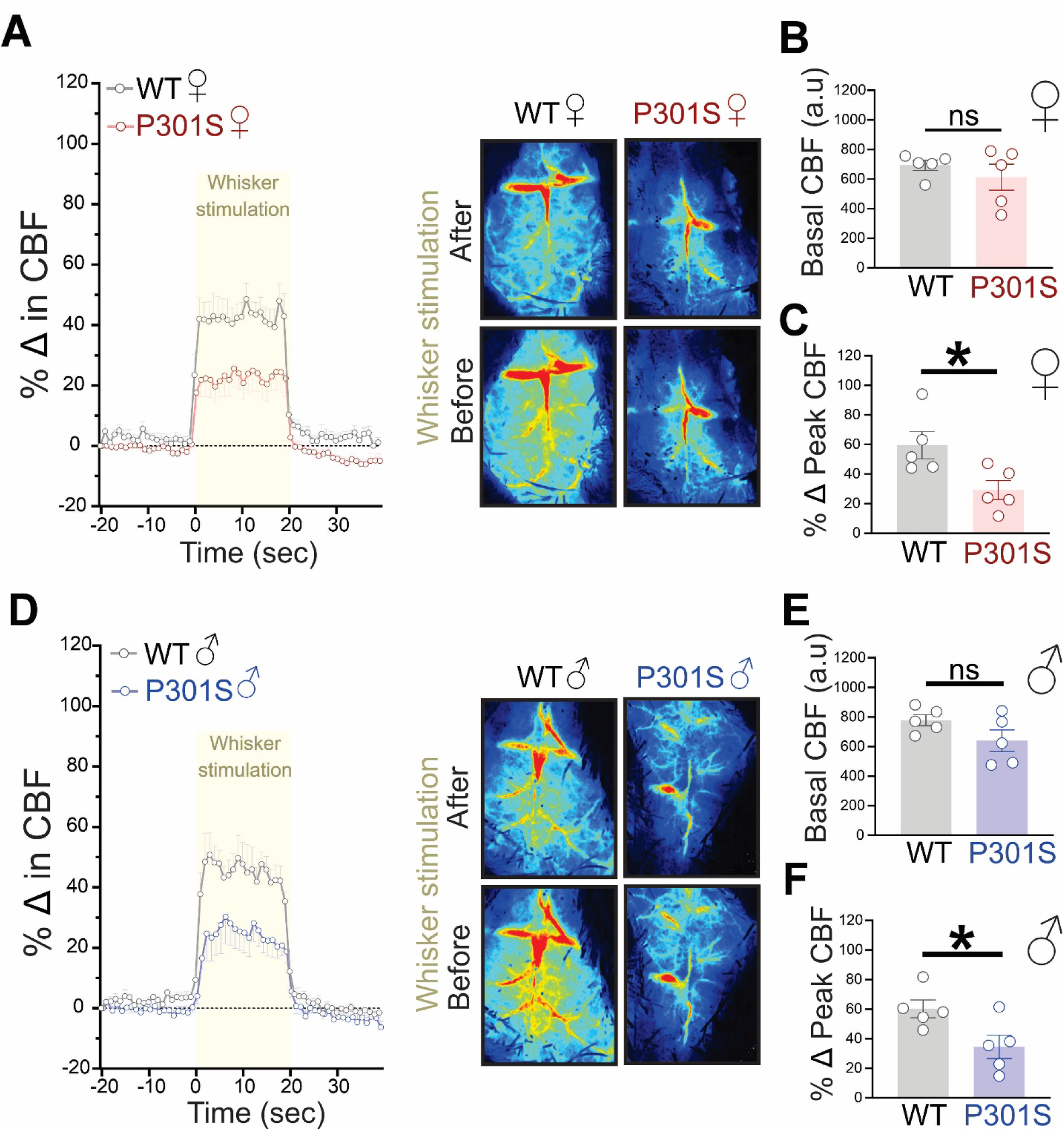
Impaired neurovascular coupling in male and female P301S tau mice. (**A**) Time course % change in cerebral blood flow (CBF) and representative laser speckle contrast images showing whisker stimulation-induced increases in CBF, (**B**) basal CBF and (**C**) peak % change in CBF upon whisker stimulation in 10-month-old female wild-type (WT) and P301S tau mice. (**D**) Time course % change in CBF and representative laser speckle contrast images showing whisker stimulation-induced increases in CBF, (**E**) basal CBF and (**F**) peak % change in CBF upon whisker stimulation in 9-10-month-old male WT and P301S tau mice. *p<0.05 by unpaired student t-test (n=5 per group). Error bars denote SEM.

### VU0486846 restores impaired neurovascular coupling in female and male P301S tau mice

To test whether M1 mAChR PAM can improve tau-mediated impairment in NVC, following baseline CBF recording, mice were injected with VU0486846 (3 mg/kg, I.P.) and changes in whisker stimulation-induced increase in CBF were reassessed every 15 min following VU0486846 treatment over 60 min period (**Fig. 3A**). Acute treatment with VU0486846 did not significantly affect whisker stimulation-induced changes in CBF in female or male wild-type mice compared to baseline (**Fig. 3B &C and 4A &B**). However, a significant increase in %Δ in CBF following whisker stimulation was observed and reached its peak at 30 min of VU0486846 treatment in female P301S tau mice (**Fig. 3D-F**), indicating that M1 mAChR PAM can restore NVC in tau mice. Similarly, in male P301S mice, VU0486846 treatment significantly increased %Δ in CBF following whisker stimulation while reaching its peak at 45 min of VU0486846 treatment (**Fig. 4C-E**), suggesting a sex-biased effect of M1 mAChR modulation where females appeared to respond earlier to VU0486846-mediated improvement in CBF. Upon comparing wild-type vs P301S tau mice responses to VU0486846 treatment, P301S tau mice showed significantly higher % increases in CBF in response to VU0486846 compared to their sex-matched wild-type controls starting at 30 min treatment in both females and males (**Fig. 3G and 4F**). Together, these results demonstrate that acute M1 mAChR positive allosteric modulation effectively rescues tau-induced NVC deficits in P301S tau mice of both sexes, with females exhibiting a more rapid response than males. Notably, the lack of an effect in wild-type mice demonstrates that VU0486846 preferentially targets disease-associated neurovascular dysfunction while sparing normal cerebrovascular regulation.

**Figure 3.**
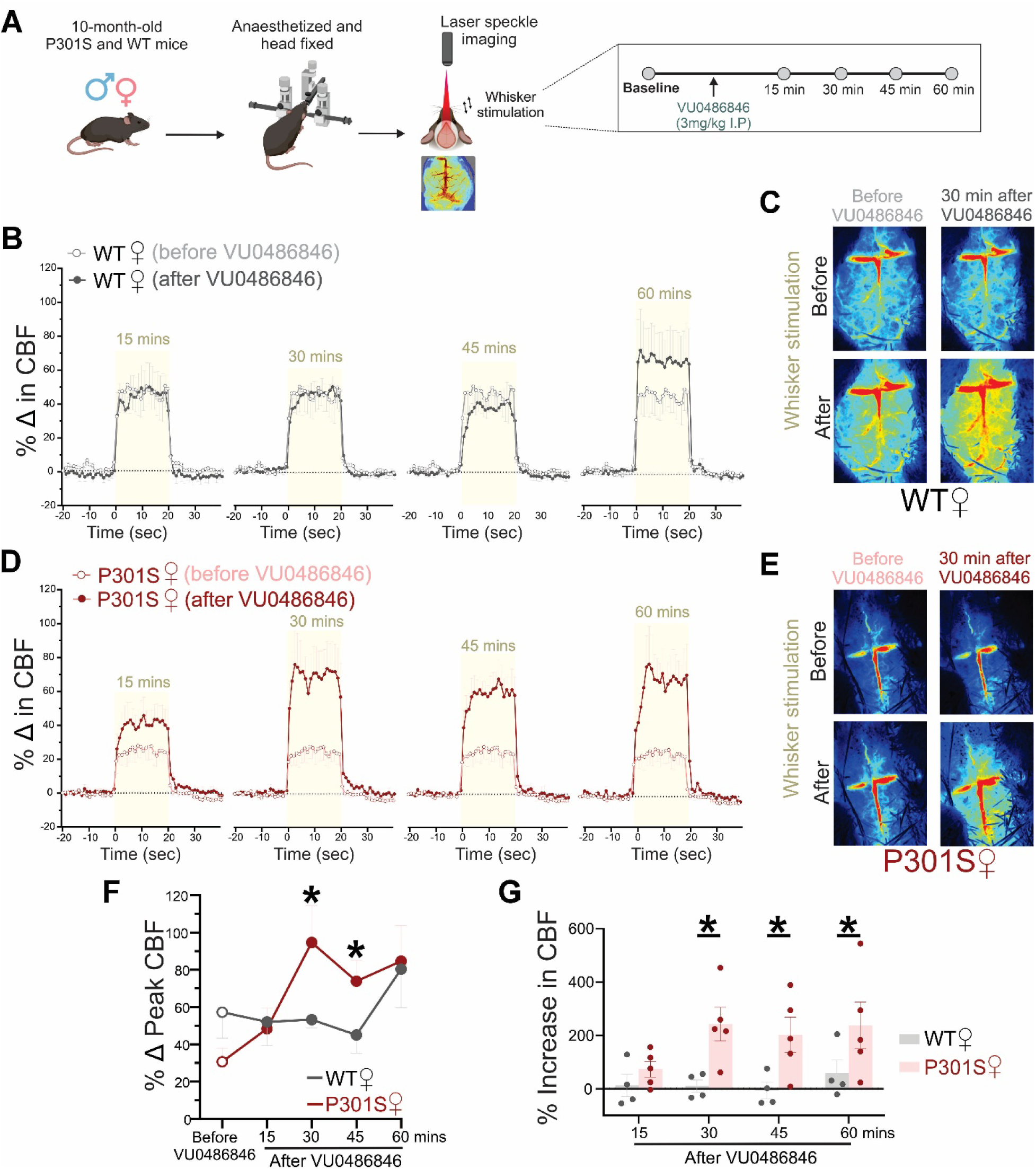
VU0486846 reverses tau-mediated impairment in neurovascular coupling in female P301S tau mice. (**A**) Schematic of the experimental design showing acute VU0486846 treatment (3 mg/kg, i.p.) followed by assessment of cerebral blood flow (CBF) using laser speckle contrast imaging in 10-month-old female and male wild-type (WT) and P301S tau mice. (**B**) Time course % change in CBF and (**C**) representative laser speckle contrast images showing whisker stimulation-induced increases in CBF in female WT mice. (**D**) Time course % change in CBF and (**E**) representative laser speckle contrast images showing whisker stimulation-induced increases in CBF in female P301S tau mice following VU0486846 treatment. (**F**) Peak % change in CBF upon whisker stimulation in female WT and P301S tau male mice following VU0486846 treatment. *p<0.05 before vs after treatment by two-way ANOVA. (**G**) % Increase in peak CBF upon whisker stimulation in female WT and P301S tau male mice following VU0486846 treatment. *p<0.05 by two-way ANOVA (n=4-5 per group). Error bars denote SEM.

**Figure 4.**
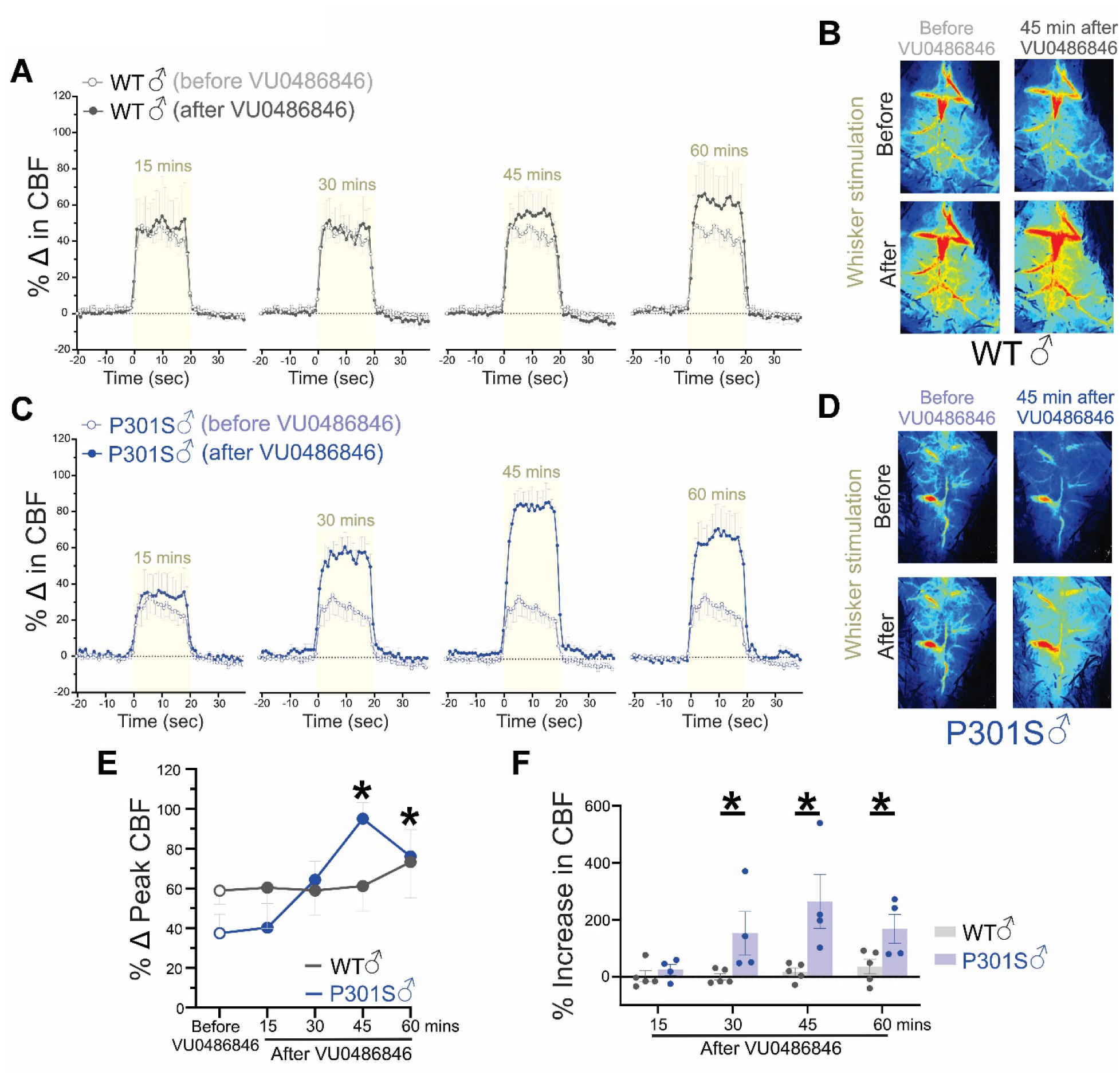
VU0486846 reverses tau-mediated impairment in neurovascular coupling in male P301S tau mice. (**A**) Time course % change in cerebral blood flow (CBF) and (**B**) representative laser speckle contrast images showing whisker stimulation-induced increases in CBF in male wild-type (WT) mice. (**C**) Time course % change in CBF and (**D**) representative laser speckle contrast images showing whisker stimulation-induced increases in CBF in male P301S tau mice following VU0486846 treatment. (**E**) Peak % change in CBF upon whisker stimulation in male WT and P301S tau male mice following VU0486846 treatment. *p<0.05 before vs after treatment by two-way ANOVA. (**F**) % Increase in peak CBF upon whisker stimulation in male WT and P301S tau male mice following VU0486846 treatment. *p<0.05 by two-way ANOVA (n=4-5 per group). Error bars denote SEM.

### Female mice exhibit faster neurovascular coupling dynamics compared to males

An interesting observation from the laser speckle analysis was that female mice appeared to reach peak CBF responses during whisker stimulation and return to baseline more rapidly than male mice. To quantify these differences in CBF dynamics, we performed a slope analysis of the stimulation-evoked responses. Linear regression was used to generate a line of best fit for the % increase in CBF from baseline to plateau, beginning at the onset of the response and ending at the plateau phase. Similarly, a second line of best fit was generated for the % decrease in CBF from plateau back to baseline, starting at the onset of the decline and ending when baseline levels were reached. The resulting slopes were used as measures of the rates of CBF increase and recovery (**Fig. 5A & C**).

**Figure 5.**
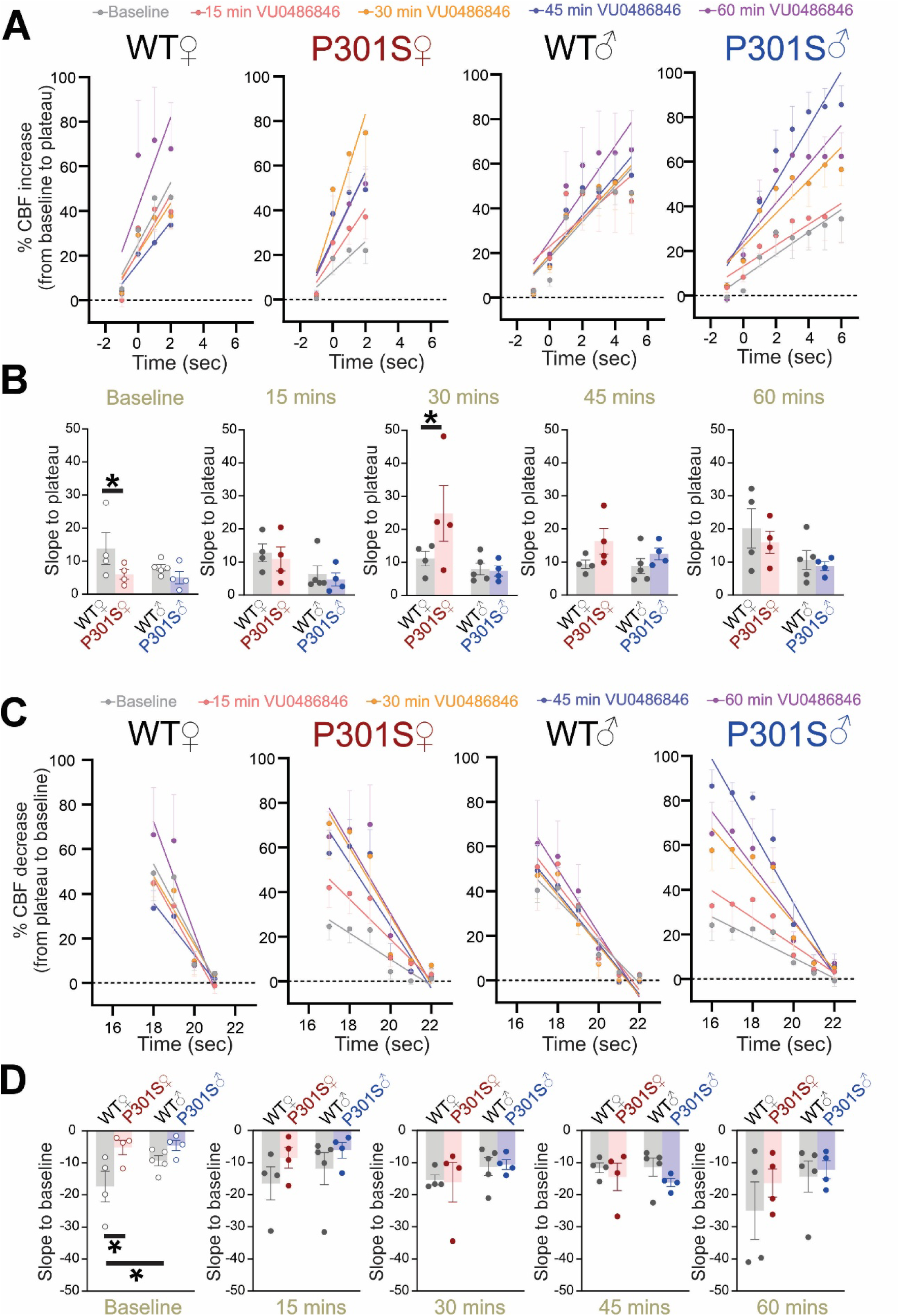
VU0486846 differentially alters the neurovascular dynamics upon whisker stimulation in female and male P301S tau mice. (**A**) Line of best fit of % increase in cerebral blood flow (CBF) from baseline to plateau and (**B**) Slope to plateau following whisker stimulation in female and male wild-type (WT) and P301S tau mice at baseline and 15, 30, 45, and 60 min after acute VU0486846 treatment (3 mg/kg, i.p.). (**C**) Line of best fit of % decrease in CBF from plateau to baseline and (**D**) slope to baseline following whisker stimulation in female and male WT and P301S tau mice at baseline and 15, 30, 45, and 60 min after acute VU0486846 treatment (3 mg/kg, i.p.). *p<0.05 by two-way ANOVA (n=4-5 per group). Error bars denote SEM.

Both wild-type and P301S tau female mice exhibited faster NVC responses than their male counterparts, requiring less time to reach peak CBF during whisker stimulation. Female wild-type mice also required less time to return to baseline following cessation of whisker stimulation (**Fig. 5A &C**). Prior to VU0486846 treatment (at baseline), only female wild-type mice displayed significantly steeper slopes for both the rise to plateau **(Fig. 5B)** and the return to baseline compared to female P301S tau mice **(Fig. 5D).** Interestingly, the slope to baseline was steeper in female wild-type mice compared to male wild-type ones (**Fig. 5D**). No significant differences were detected between male wild-type and P301S tau mice in the rise-to-plateau or return-to-baseline slopes (**Fig. 5B &D**). These findings indicate that neurovascular responses are more robust in female mice than in males and that tau pathology disproportionately impairs this enhanced neurovascular responsiveness.

Following VU0486846 treatment, no significant changes in slopes were detected between male wild-type and P301S tau mice. In contrast, the baseline differences observed between female wild-type and P301S tau mice were abolished as early as 15 min following treatment. By 30 min post-VU0486846 treatment, female P301S tau mice exhibited a significantly steeper slope to plateau than female wild-type controls (**Fig. 5B &D**). Overall, these results identify sex-specific differences in neurovascular dynamics and demonstrate that tau pathology preferentially disrupts the rapid cerebrovascular responses observed in females. Additionally, they establish M1 mAChR activation as a promising strategy for restoring neurovascular function and overcoming tau-induced neurovascular dysfunction.

## Discussion

Reduced CBF is increasingly recognized as an early contributor to Alzheimer’s disease, occurring years before the onset of cognitive decline.^25^ Rather than being just a consequence of neurodegeneration, chronic cerebral hypoperfusion and impaired NVC are now considered major contributors to the disease progression.^26^ Importantly, adequate cerebral perfusion is essential for the clearance of toxic proteins, including β-amyloid and hyperphosphorylated tau. Consequently, sustained reductions in CBF can compromise β-amyloid and tau clearance, accelerating synaptic dysfunction and cognitive decline.^27^ Although restoring cerebral perfusion and NVC has emerged as an attractive therapeutic strategy, clinical translation has been limited by our incomplete understanding of the signaling pathways and the specific receptors involved in these processes. We demonstrate that M1 mAChR activation restores NVC in mice with Alzheimer’s disease, uncovering a previously unrecognized neurovascular mechanism underlying the therapeutic benefits of M1 mAChR modulation. Although both sexes benefited from treatment, the earlier neurovascular recovery observed in female mice suggests that M1 mAChR signaling regulates neurovascular dynamics in a sex-dependent manner. These findings highlight a strong association between tau pathology, cognitive decline and neurovascular dysfunction during Alzheimer’s disease, and provide further evidence that M1 mAChR could be a promising disease-modifying therapeutic target for Alzheimer’s disease via modulation of NVC.

Tau hyperphosphorylation during Alzheimer’s disease is known to disrupt neuronal function contributing to the disease neurotoxic insult, which is basically associated with impaired cognitive function occurring at the early stages of Alzheimer’s disease.^28^ This is consistent with our findings showing that P301S tau mice exhibited significant deficits in learning and memory, particularly recognition and spatial memory functions as evidenced by the novel object recognition and Morris water maze tests respectively, further supporting the validity of P301S mice as a model of Alzheimer’s disease -related pathology.

Additionally, neurovascular function was significantly impaired in P301S tau mice evidenced by the marked reduction in whisker stimulation-induced increase in CBF in both male and female P301S tau mice compared to wild-type controls, reflecting impaired NVC, which goes in line with the noticeable cognitive decline observed in P301S tau mice. This is also consistent with previous reports demonstrating that Alzheimer’s disease and tauopathy are associated with neurovascular dysfunction.^29–31^ Interestingly, recent studies in PS19 tau mouse model reported that tau disrupts NVC even before the formation of neurofibrillary tangles and the development of cognitive decline, demonstrating impaired NVC as an early pathological event during Alzheimer’s disease that may accelerate tau-mediated pathology.^32^ Together with our findings, this supports the notion that impaired NVC in P301S mice plays critical roles in Alzheimer’s disease pathophysiology and its associated neurodegeneration.

The cholinergic system is a key regulator of both cognitive function and CBF, and impaired activity of M1 mAChR is a hallmark of Alzheimer’s disease. M1 mAChR is highly expressed in different brain regions critical for memory and learning.^33, 34^ M1 mAChR signaling also contributes to NVC regulation, as cholinergic activation can induce vasodilation via nitric oxide-dependent mechanisms and by interacting with endothelial cells.^35, 36^ Impaired cholinergic signaling has been reported in different animal models of Alzheimer’s disease and enhancing it using acetylcholinesterase inhibitors such as donepezil has been shown to increase CBF and neuronal functional connectivity reflected on cognitive performance.^15, 16^ Our group has also shown that the use of M1 mAChR PAM can rescue cognitive function in both male and female mice with Alzheimer’s disease.^8, 11^ In line with this, our present study shows that acute treatment with VU0486846 resulted in a robust and rapid improvement in whisker stimulation-induced increase in CBF in both male and female P301S tau mice, underscoring improved NVC response. This effect peaked between 30- and 45-min post-treatment, suggesting a rapid onset of action. Importantly, VU0486846 had no significant effect on whisker stimulation-induced increase in CBF in wild-type mice, indicating that its action is dependent on the presence of underlying pathology. Altogether, this underscores the ability of M1 mAChR modulation to restore impaired signaling pathways rather than enhancing normal physiological function and support the concept that restoring cholinergic function may alleviate neurovascular abnormalities, a key driver of Alzheimer’s disease progression. The mechanisms underlying this improvement in NVC may involve increased nitric oxide production and vasodilation in response to M1 mAChR activation, restoring communication between neurons and vascular cells, or reducing neuroinflammation and glial cells activation, which are known contributors to vascular dysfunction and the loss of blood brain barrier integrity during Alzheimer’s disease as previously reported.^35–38^

Considering the well-documented sex differences in Alzheimer’s disease pathology and prevalence,^39^ a notable finding of the study is the presence of sex-specific response to VU0486846. While both male and female P301S tau mice exhibited significant improvements in NVC response, female mice showed an earlier peak response at 30 min, compared to 45 min in males. Notably, a study of post-mortem brain tissue from Alzheimer’s disease patients identified sex-dependent differences in M1 mAChR expression in the temporal cortex, suggesting that cholinergic signaling may be differentially regulated between females and males in brain regions involved in learning and memory.^40^ In addition, estrogen is a key modulator of both cholinergic signaling and cerebrovascular function, enhancing M1 mAChR signaling while promoting vasodilation through endothelial nitric oxide pathways.^41–43^ Together, these observations provide a potential biological basis for the sex-dependent differences in neurovascular dynamics observed in the present study and further highlight the importance of considering sex as a biological variable in the development of M1 mAChR-targeted therapies for Alzheimer’s disease.

One of the intriguing findings of our study is the identification of sex-dependent differences in neurovascular dynamics under physiological conditions. Female wild-type mice exhibited faster NVC responses than males, reflected by a more rapid return of CBF to baseline following whisker stimulation. Strikingly, this advantage was completely lost in P301S tau mice, suggesting that tau pathology selectively disrupts the enhanced neurovascular responsiveness present in healthy females. Importantly, M1 mAChR activation restored these neurovascular dynamics, revealing a previously unrecognized role for M1 mAChR signaling in regulating the temporal properties of NVC during tau pathology. These findings not only identify neurovascular dynamics as a sensitive readout of tau-induced dysfunction but also highlight M1 mAChR activation as a promising strategy to restore neurovascular function in Alzheimer’s disease, with potential sex-dependent differences in therapeutic responsiveness.

While our findings establish a previously unrecognized role for M1 mAChR activation in restoring neurovascular function, our findings should be interpreted within the limitations of the current study. Future studies are warranted to determine the long-term therapeutic benefits of M1 mAChR positive allosteric modulation on NVC and its impact on disease progression, as well as to further define the cellular and molecular mechanisms by which M1 mAChR signaling preserves neurovascular integrity in Alzheimer’s disease.

In conclusion, our findings uncover a novel role for M1 mAChR signaling in maintaining neurovascular integrity during tau pathology. By demonstrating that M1 mAChR activation restores NVC and CBF, we identify preservation of neurovascular function as a previously unrecognized disease-modifying mechanism underlying the therapeutic benefits of M1 mAChR activation in Alzheimer’s disease. These findings broaden the therapeutic potential of M1 mAChR-targeted therapies beyond their established cognitive benefits and position M1 mAChR as a promising target for simultaneously restoring neuronal and neurovascular function in Alzheimer’s disease.

## Data availability

The data underlying this article is available by the corresponding author upon reasonable request.

## Authors contributions

**Wesam Bassiouni:** Conceptualization, Investigation, Data Curation, Analysis, Writing-Original Draft. **Mohamed Abdelnaby:** Investigation, Data Curation, Analysis. **En-Hung Ai:** Investigation, Data Curation, Analysis. **Khaled S. Abd-Elrahman:** Conceptualization, Investigation, Analysis, Funding Acquisition, Resources, Supervision and Project Administration, Writing-Review and Editing.

## List of Abbreviations

CBF: Cerebral blood flow
M1 mAChR: M1 muscarinic acetylcholine receptor
NVC: Neurovascular coupling
PAM: Positive allosteric modulator

